# Single cell analysis reveals dynamics of transposable element transcription following epigenetic de-repression

**DOI:** 10.1101/462853

**Authors:** David Brocks, Elad Chomsky, Zohar Mukamel, Aviezer Lifshitz, Amos Tanay

**Affiliations:** Weizmann Institute of Science, Department of Applied Mathematics and Computer Science and Department of Biological Regulation

## Abstract

Spontaneous or pharmacological loss of epigenetic repression exposes thousands of promoters encoded in transposable elements (TEs) for pervasive transcriptional activation. How TE responses differ between epigenetically relaxed cancer cells and what factors govern such variation remains however largely unknown. By quantifying TE transcription initiation at single cell and locus resolution in epigenetically targeted cancer cells, we characterize specific groups of co-regulated loci that drive over ten-fold variation in TE load per single cell. Such variable activity patterns are largely linked to cell cycle stages, stress response signatures, and immune pathways. Furthermore, cells with high levels of specific transcription factors show increased TE expression, while within such cells, multi-copy families are differentially regulated in response to local sequence divergence of binding sites and the locus’ repressive or active chromosomal contexts. Our data thereby implicates the regulation of potent promoters within TEs as an underestimated source of transcriptional heterogeneity following epigenetic therapy.

## INTRODUCTION

Transposable elements (TEs) are DNA sequences that multiply in the germline via vertical transmission. They have been evolutionary successful in colonizing many eukaryotic genomes and their massive expansion lead to partial neo-functionalization, in particular as regulatory elements^1,2^. Inactivating mutations have neutralized the threat of insertional mutagenesis for most human TEs. Nevertheless, retrotransposons (LINEs, SINEs, and LTRs) that require an RNA intermediate for their life-cycle encode strong promoter activity that poses the risk of spurious transcriptional activation and subsequent epigenetic perturbation across numerous genomic loci^3,4^. Host cells therefore utilize multiple epigenetic surveillance mechanisms, such as DNA methylation and histone modifications, to prevent the pervasive mis-expression of TEs^5,6^.

Accordingly, pharmacological targeting of any such suppressive mechanisms may provoke immune responses in cancer cells by releasing TEs as a source for neoantigens or virus-mimicking double-stranded RNAs^7,8^. Since the initial discovery of this effect for DNA hypomethylating agents, the list of drugs triggering ‘viral mimicry’ has been steadily extended and now even includes molecules with no direct target in the epigenetic enzyme machinery^9,10^. The surprisingly broad spectrum of TE activation by distinct classes of anti-cancer drugs illustrates our lack of knowledge about the factors governing their derepression propensity. For example, while TEs are targeted by common mechanisms for epigenetic repression, their ancient promoters are diverse at multiple levels. TE sequences are originating from many classes and families, which continue to diverge even after their immobilization, creating diverse repertoires of cis-elements that can promote binding by different trans-factors^11,12^. The genomic and epigenomic contexts of TEs are also diverse, with specific families enriched in repressed or active genomic domains, within proximity to other regulatory elements, or away from them^13^. How these factors contribute to the potentially complex and locus-specific regulation of TEs, especially once epigenetic repression is eroded or targeted therapeutically remains unknown^7,8,14^.

Here, we develop a 5’ single-cell RNA sequencing (scRNA-seq) strategy with integrated TE and genic analysis pipeline, to map active transcription start sites (TSSs) within TEs *de novo* and model their co-regulation within sub-populations of single cells. We applied our new approach to study two cancer cell lines following treatment with a clinically established DNA hypomethylating agent and HDAC inhibitor, a drug combination known to induce massive expression of specific TE families^14^. Unexpectedly, we found highly heterogeneous overall TE activity levels in different populations of single cells. This heterogeneity is driven by specific groups of co-regulated TEs, which in turn were strongly linked with fundamental biological processes, including the cell cycle, stress responses, and type-I interferon signaling. Activity patterns of loci from genetically similar families were correlated with specific sequence polymorphisms in their putative promoters, the expression of corresponding transcription factors (TFs), as well as with the transcriptional permissiveness of their genomic neighborhood. 5’ scRNA-seq therefore highlights the necessity to deconvolute TEs’ transcriptional output to the single cell and locus level. Using this new technology it is now possible to investigate poorly appreciated aspects of genome regulation in epigenetically de-repressed cells, not only following epigenetic therapy, but during specific embryonic stages^15^, aging^16^, and carcinogenesis^17^.

## RESULTS

### Epigenetic therapy induces heterogeneous TE de-repression in single cancer cells

We cultured lung and colon cancer cell lines (H1299 and HCT116, respectively) and performed 5’ scRNA-seq to map the transcriptional response to an established epigenetic drug regimen combining a low-dose DNA hypomethylating agent and an HDAC inhibitor (DACSB). After exclusion of low-quality cell barcodes (**Fig S1A**), we retained over 15,000 single cell profiles from treated and untreated cells, with a median gene transcript count of 7972 to 17328 unique molecular identifiers (UMIs) per cell across conditions (**Fig 1A**). We further confirmed UMI enrichment towards the 5’ start of gene models, allowing us to de novo identify between 11,448 and 14,756 putative TSS with high consistency between conditions (**Fig S1B-F**). The resulting TSS intervals accounted for ∼62% of the total genic signal and the remaining UMIs could be attributed to transcriptional noise or technical effects (**Fig S1G-H**).

**Figure 1:**
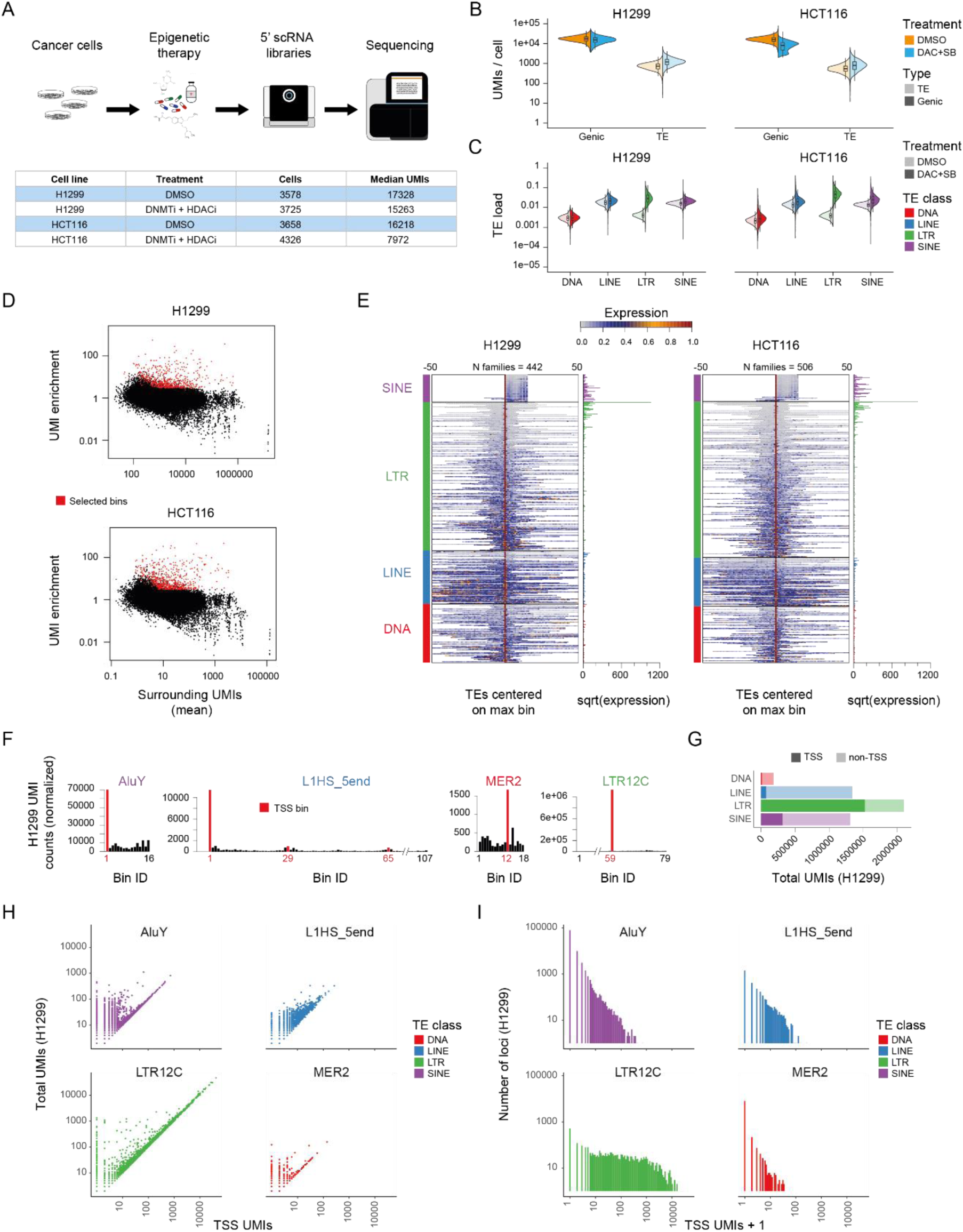
Single-cell RNA-seq identifies the transcription start site of genes. A) Summary of the applied experimental workflow. B) Violin plot showing the single-cell density distribution of total genic and TE UMI counts for untreated (red) and treated (blue) H1299 (left) and HCT116 (right) cells. C) Single-cell density distribution of TE load (TE UMIs / total UMIs) in four classes, before (light colors) and after (dark colors) treatment. D) Enrichment scores for TE consensus bins (Y axis), by total UMI counts on neighboring bins for H1299 (top) and HCT116 (bottom) cells. Bins identified as putative TSSs are highlighted in red. E) Aggregate normalized UMI count distribution along the consensus models for families with at least 1 detected TSS bin in H1299 (n=442, left) and HCT116 (n=506, right) cells. Family profiles (rows) are centered around the TSS bin with highest raw expression and arranged by TE class (colored boxes) and TSS bin enrichment. UMI aggregate counts of the maximum TSS bin are shown in the right. F) Distribution of normalized UMI counts along the consensus model of selected (highest overall expression in H1299) representatives from each TE class. Putative TSS bins are highlighted in red. G) Number of TSS and non-TSS UMIs mapping to different TE classes in treated H1299 cells. H) Per locus total number of UMIs as a function of TSS-only UMIs for representative TE families. For each TE class, the maximally expressed family in H1299 cells was selected. I) Distribution of total UMIs per locus for representative TE families.

We also identified molecules mapping unambiguously to individual TE loci, and confirmed that the treatment triggered on average a 1.56 – 1.72 (H1299 and HCT116) fold transcriptional induction of TEs in both cell lines (**Fig 1B**). This allowed us to quantify TE transcription at single cell resolution with high accuracy based on 19 – 7037 (12 – 6444 for HCT116) TE UMIs per cell post treatment. Interestingly, both cell lines showed profound intercellular heterogeneity in their TE transcriptional load (7.53 and 7.87-fold change between 5 and 95 percentile in H1299 and HCT116, respectively), and in particular in LTR activity (10.5 and 9.35 fold variation, **Fig 1C**). The surprising extent of global TE expression heterogeneity motivated further in-depth analysis of locus- and family-specific variability in TE de-repression, and the factors contributing to it.

### 5’ scRNA-seq defines putative TSSs in thousands of TE loci

Encouraged by the robust identification of genic TSSs based on 5’ scRNA-seq data, we adapted our approach to define transcriptional activity originating in TEs. We focused solely on UMIs that uniquely mapped to individual genomic TE loci (between 86.17 and 88.96% of all TE UMIs post treatment, **Fig S2A**), which allowed the unbiased quantification of transcriptional activity for most families (**Fig S2B**). To improve power in detecting bona-fide TSS activity within TEs, we initially focused on treated cells and aggregated their UMIs over consensus sequence models for 1076 families (between 3 to 228,527 loci per model) representing the four major TE classes (LTR, LINE, SINE, and DNA) (**Fig 1D)**. We binned the consensus sequence model of each family and analyzed relative enrichment of bins as a proxy for specific TSS activity (**Fig 1E**). This led to the identification of 722 bins in 442 families with putative TSS activity in H1299 cells (862/506 in HCT116). As shown in **Fig 1F**, mapping UMIs on the TE consensus model for families with at least one specific bin reflected a highly specific pattern of localized consensus transcription initiation (see **Fig 1G and Fig S2C** for case examples). We validated that this effect is not correlated with variable mapping efficiency in degenerate TE sequences (**Fig S2D**). Although the degree of TSS specificity was markedly different among broad classes of TEs **(Fig S2E**), the overall transcriptional density (molecules per kb), in non-TSS TE bins was comparable (**Fig S2F)**. This showed elevation in transcriptional output was indeed specific for TSSs in certain families rather than a consequence of broad patterns of genomic de-repression without promoter specificity. Applying the same approach to untreated cells, we found little TE signal originating from bona-fide TSS activity (**Fig S2G**), which is why we focused downstream analyses on epigenetically de-repressed cells. In conclusion, we observed specific transcriptional initiation in consensus models of de-repressed TE families (and LTRs in particular) following treatment which facilitates subsequent inference of activity for individual TE copies at single cell resolution.

To better understand potential intra-TE family expression dynamics, we decomposed consensus TSS signals back into their single components and investigated the contribution of individual TE copies to the TE consensus signal. We found that most of the active families aggregated transcriptional contribution from a large number of expressed loci (329 out of 442 expressed families with over 10 contributing loci in H1299, 405 out of 507 in HCT116) (**Fig S3A**). For highly expressed families, we also confirmed that the specificity of the TSS bin was completely consistent between copies (**Fig 1H and Fig S3B**). On the other hand, for all TE families, the transcriptional response was incomplete and a large number of TE loci remained repressed (**Fig 1I and Fig S3C**). For the most consistently active family (LTR12C), we observed transcriptional activity in 2208/2310 (in H1299/HCT116) out of 2735 loci. Although half of all LTR12C UMIs were contributed by only 3.17/3.38% (n=70/78) of the loci, intrafamily sequence diversity allowed us to identify true transcriptional activity for the remaining copies rather than counting misaligned UMIs from few high output TSSs **(Fig S3D and S3E**). To validate the inferred TE expression patterns, we confirmed a high degree of consistency in the activity of TE loci between the two treated cancer cell lines (**Fig S3F**), in particular for loci associated with high expression. In conclusion, we defined 136,366 genomic loci from 442 TE families (170,746 from 506 in HCT1160) as sites of potential transcriptional initiation following epigenetic therapy. The activity in these sites generated 0 – 23.9% (0 – 38% HCT116) of the total RNA molecule count profiled per cell (3.1 and 5.85% on average, respectively), suggesting a considerable influence on the genomic landscape of transcriptional initiation which goes beyond canonical assembly of transcriptional machineries at genic TSSs. This highlights the importance of understanding the mechanisms driving the specification and activation of TE TSSs at single cell and single locus resolution.

### TE metacells uncover co-regulatory TE modules

Having identified transcriptional output from thousands of TE loci at single-cell resolution, we next searched for cell subpopulations that share TE expression patterns, as a way to define TE co-regulatory modules. Analysis of normalized single cell transcriptional variance identified high variance TE TSS loci in treated cells (**Fig 2A**). Those TSSs also showed a detailed and non-homogeneous correlation structure (**Fig 2B** and **Fig S3G**) within the single cell cohort, confirming their expression is not a mere reflection of variable global TE load following treatment. Furthermore, TE clusters that were correlated in their expression patterns also showed a high degree of pairwise sequence similarity and often mapped to phylogenetically related families (**Fig 2B**, lower right). We organized correlated TEs into groups and annotated the derived TE modules (10 for each cell line) based on their dominant TE class/superfamily associations. This resulted in the identification of multiple ERV modules (comprising mostly solitary LTRs) with a complex correlation structure, a HCT116-specific LINE/DNA module, as well as 2 modules (3 in HCT116) defined by weaker association of multiple Alu elements.

**Figure 2:**
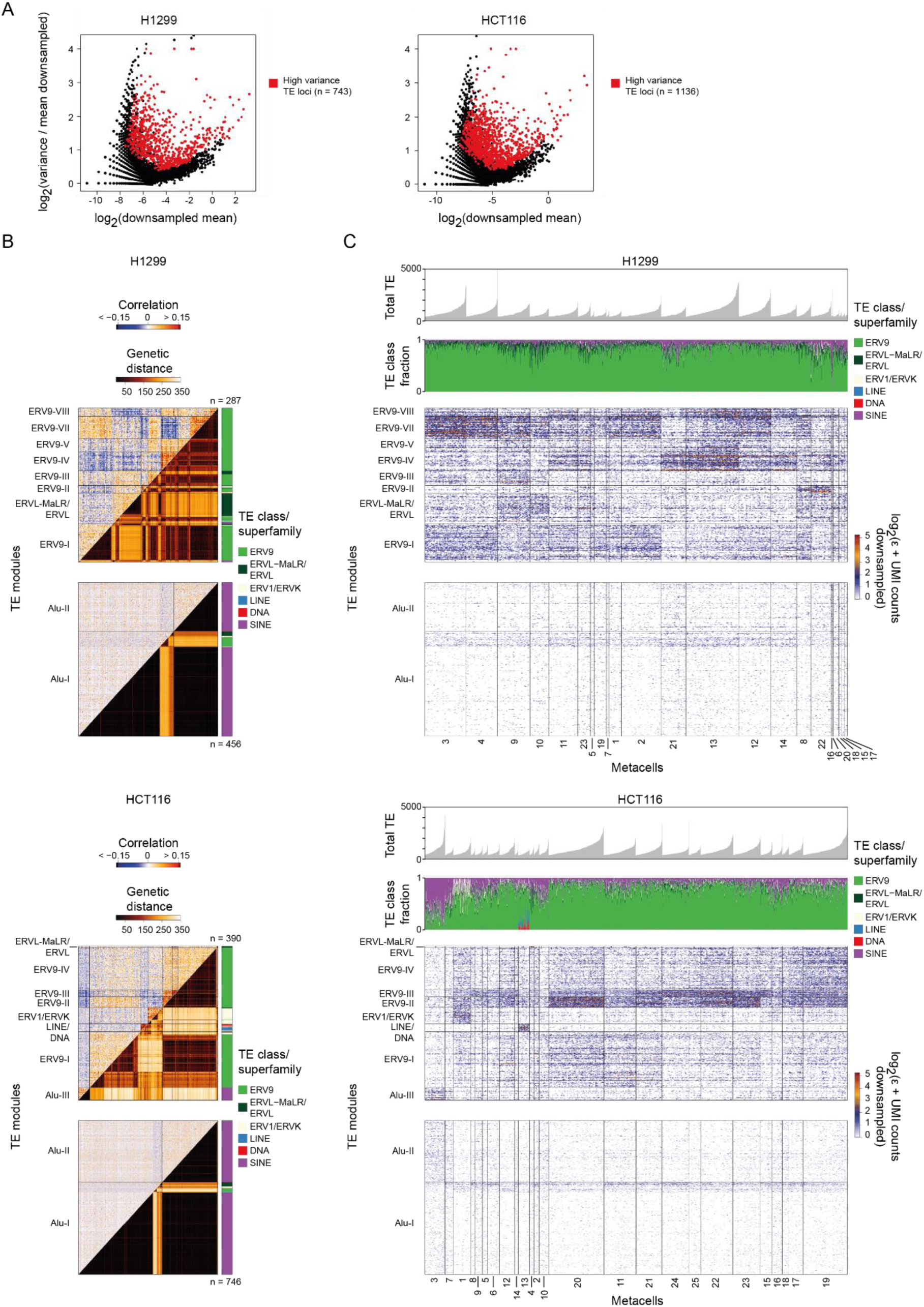
Meta-grouping of cells with consistent TE expression identifies structured variation in TE transcription. A) Normalized variance vs. mean UMI counts in down-sampled TE TSSs UMI matrices. B) Single-cell expression correlation (upper triangle) and pairwise genetic divergence (lower triangle) for 10 TE TSSs modules in H1299 (top) and HCT116 (bottom) cells. SINE and non-SINE modules are plotted separately for clarity. Color annotation represent a loci’s TE class or superfamily membership. C) H1299 (upper) and HCT116 (lower) single-cell expression matrix showing the rolling mean (five cells) of downsampled and log2 transformed UMI counts. Rows represent individual loci arranged by TE modules and columns correspond to individual cells grouped by metacells and arranged by total TE load. The two upper bar plots indicate a cell’s absolute TE load and the contribution of each TE class to it.

We adapted the MetaCell framework^18^ to identify TE metacells as groups of single cells showing highly similar TE co-expression patterns. Analysis of TE metacells and the distribution of TE expression over them showed that the rich correlation structure among TE TSSs is reflecting an organization of the single cell cohort into groups representing distinctive combinatorial expression signatures (**Fig 2C**). Importantly, although the observed metacells differed considerably in their overall TE burden (**Fig 2C**, top panel), some of the co-expressed TE TSS modules behaved antagonistically. This was the case even for modules grouping together the same superfamily of ERV elements (e.g. modules ERV9-IV and ERV9-VII in H1299) (**Fig S3G**), confirming metacells were identifying combinatorial TE regulation rather than approximating different degrees of overall TE activity. These findings demonstrate regulated TE expression dynamics in treated cells and suggest the presence of inter- and intra-family specific transcriptional regulation going beyond global and stochastic epigenetic de-repression.

### Gene expression correlates with specific TE activation signatures

The observed TE subpopulation dynamics hint towards coordinated control of TE modules by factors acting in trans. In that case, TE metacells, which are statistically derived based solely on TE expression, should be distinguishable also by their gene expression signatures. Consistent with this idea, we observed remarkably rich gene expression signatures in TE metacells (**Fig 3A-C**), that exceeded any localized effect of TE de-repression on nearby genes (**Fig S4A**). Globally, we detected 6518 (3950 in HCT116) (chi-squared, FDR < 0.01) genes with significant differential expression between TE metacells. More specifically, genes enriched in specific TE metacells were functionally and spatially related. For example, in H1299 we observed KRT8, KRT18 and additional genes overexpressed in TE metacells #22 and #8, expressing TE module ERV9-II (**Fig S4B**). In HCT116, expression of early-immediate genes (JUN, ARC) were enriched in TE metacells #1 expressing TE module ERV1/ERVK (**Fig S4C**). The LINE/DNA module was highly specific for HCT116 metacell #13, comprising presumably apoptotic cells based on their high mitochondrial and autophagy-related gene expression signature (ATG10, DNASE1) (**Fig S4D**). Importantly, we identified a group of TE metacells in H1299 (#12-14, #21) that were enriched for expression of loci within chromosomal locus 19q3 (**Fig S4E and F**), suggesting potential copy number heterogeneity as the underlying mechanism for up-regulation of TEs and genes in these metacells. However, most other TEs were not linked with specific chromosomal domains, and included a balanced mixture of chromosomal loci that are unlikely to be explained by sub-clonal structure bearing specific chromosomal aberrations (**Fig S4G and H**).

**Figure 3:**
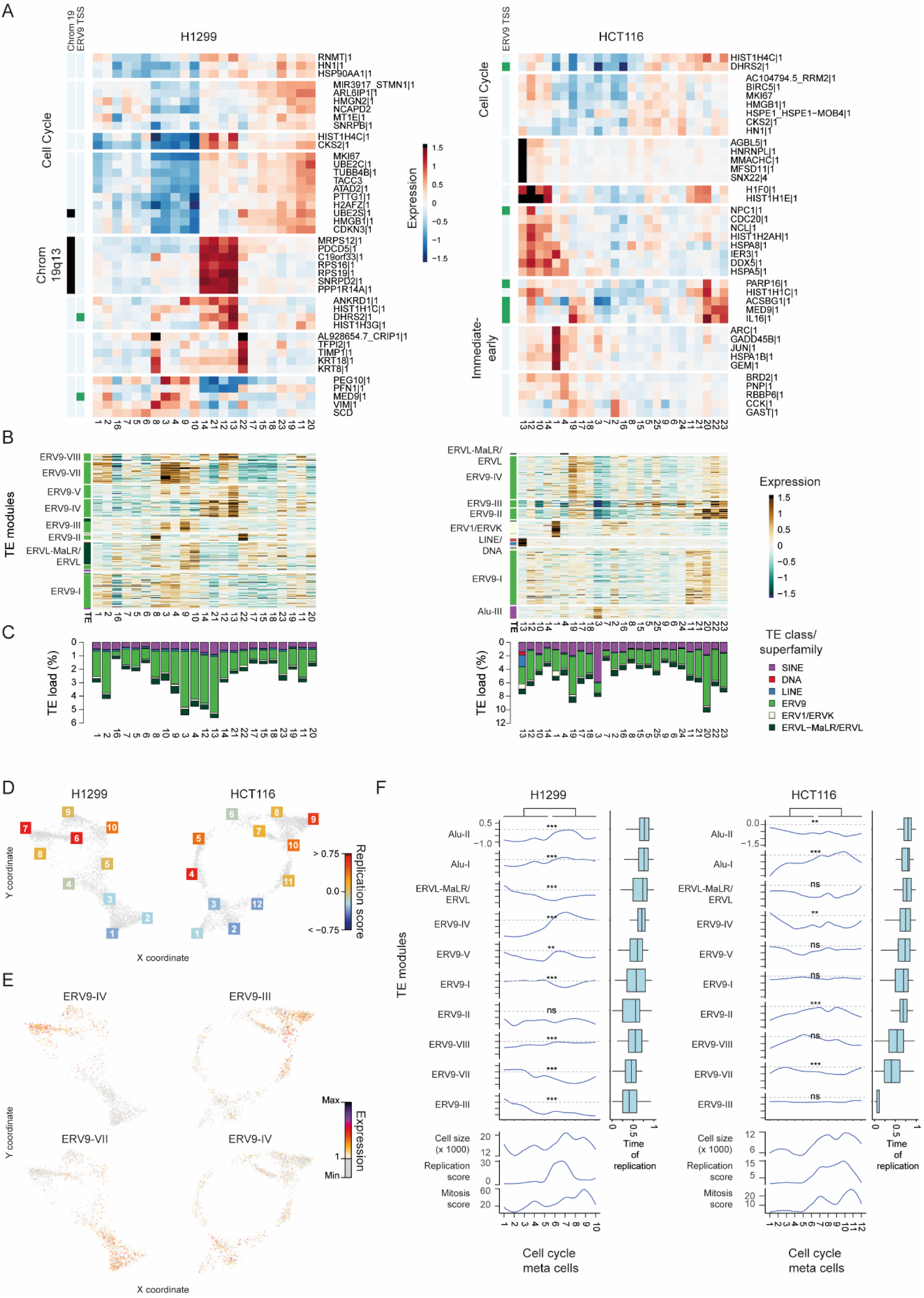
TE meta-cells are linked to the cell cycle and other gene-modules. A) Shown are enrichment values for genes (rows) over TE metacells (columns). Left color bars indicate genic TSS overlap with ERV9 family members (green) and mapping to chromosome 19 in H1299. Metacell numbers (bottom) are consistent with Figure 3. B) Enrichment values for TE loci (rows) across metacells (columns). Rows are grouped by module (as in Fig 3). Colored bars represent the superfamily/class membership of a locus. Alu modules are not shown. C) Average TE load per metacell. Colors indicate the contribution of individual TE classes to the total. D) Metacell regularized force-directed single-cell 2D projection based on cell cycle transcriptome signatures. Grey dots represent individual cells. The color of the corresponding metacell ID (squares positioned at the centroid) is based on the average expression footprint of presumed mitosis-specific genes (top 10 genes with highest footprint in mitotic metacell). E) Projection as before but colors indicate summarized expression intensity of indicated TE modules. F) Cell cycle metacells (columns) were arranged according to their estimated cell cycle phase from left (G0/G1) to right (mitotic exit) in a supervised fashion. Upper rows show the smoothed enrichment of aggregated TE modules across metacells. TE modules were arranged by their median time of replication (right boxplots). Mitosis and replication expression scores (lower rows) summarize downsampled UMI counts of the top 10 genes in the respective cell cycle gene module. Significance is based on Wilcoxon rank-sum test comparing cells from metacells #1-5 (#1-6 in HCT116) against #6-10 (#7-12 in HCT116).

To complement the detection of these combinatorial gene expression patterns in TE metacells, we searched for gene expression signatures that correlated with overall TE load in single cell resolution (**Fig S5A**). We found various immune-related genes (e.g. IL16) to be positively associated with overall TE load, in line with recent reports on de-repressed endogenous retroviruses (ERVs) triggering a viral-like interferon (IFN)- response^7,8,19^. Indeed, overall IFN-I intensity was correlated with total TE load (**Fig S5B**) and this effect was mostly driven by specific LTRs families at the metacell level (**Fig S5C**). However, the effect size was modest with at most 1.61 and 1.5 fold change in IFN-I expression between TE metacells in H1299 and HCT116, respectively. In summary, our analysis defines a rich subpopulation structure in treated cells, involving combinatorial activity of different TE TSSs, and correlated patterns of regulated gene expression. These data suggest a cancer cell’s state may regulate classical gene and TE promoters simultaneously.

### TE activation patterns are cell-cycle dependent

Some of the most notable gene expression signatures observed for TE metacells involved co-regulation of multiple cell cycle genes (e.g. MKI67, UBE2C, PTTG1). To follow up on these observations and characterize systematically potential cell cycle regulation of TE expression, we generated cell cycle metacell models using a selected set of genes in each cell line (**Fig 3D and S6A**). To our surprise, expression of different TE families (especially SINEs) largely varied between cycling and non-cycling cells, even before treatment (**Fig S6B**). Studying this effect at higher resolution, we projected the total expression of TE modules on the inferred cell cycle trajectory (**Fig 3E-F**). This resulted in observation of unexpectedly pervasive cell cycle correlation of TE expression patterns. In H1299, some TE modules (ERVL-MaLR/ERVL, ERV9-III, and ERV9-VII) peaked in expression at G1/G0, while other TE modules (ERV9-IV, and ERV9-VI) showed positive correlation of their expression with specific stages in S phase. Analogous dynamics (albeit less specific) were inferred for HCT116 cells. Interestingly, analysis of the distributions of genomic time of replication for loci in different TE modules showed remarkable diversity as well. We observed an overall enrichment of active Alu elements at early replicating domains consistent with the general preference of SINEs to active genomic loci. More surprisingly, we detected distinct time of replication distributions for different ERV9 TE modules, for example showing in H1299 the ERV9-III module as early replicating and the ERV-IV module as late replicating. The correlation between TE cell cycle dynamics and time of replication was imperfect – although the two modules with the latest time of replication in H1299 (ERV-III, ERV-VII) showed decrease in expression during replication.

Interestingly, we observed mild cell cycle correlations for some of the identified TE modules even in untreated cells, suggesting this effect is generally applicable, even when TEs are only weakly and sporadically de-repressed (**Fig S6C**). Together, these data indicate that chromosomal dynamics during replication can act together with different cellular programs to regulate specific patterns of TE activity.

### Trans and cis factors shape the complex de-repression landscape of multi-copy families

The analysis of TE metacells and TE TSS modules suggested that some of the derepression dynamics are correlated with the activity of specific trans-factors. However, since even highly responsive TE families were characterized by only partial de-repression of the family’s loci, we next aimed to define sequence features and genomics contexts that predispose loci to become de-repressed. Focusing on families contributing at least 5 loci to TE modules, we first found strong correlation between the distance of a TE locus to the nearest expressed genic TSS, and its de-repression predisposition (**Fig 4A** and **Fig S7A**). This effect was linked with enriched de-repression probability of loci in the active chromosome compartment, and lower de-repression in loci within lamina associated domains (**Fig 4B** and **Fig S7B**). Analysis of DNA methylation data before and after treatment^14^ showed that de-repressed TE loci are significantly more methylated before treatment than loci that are not de-repressed (**Fig 4C**). After treatment, TE methylation levels are decreasing around the responsive TEs, converging to levels similar to those initially observed in the non-responsive loci prior to treatment. This result is suggesting the non-responsive TE loci are relying on mechanisms other than DNA methylation for repression, while the de-repressed loci show tight correlation, and likely a causal link between methylation and repression.

**Figure 4:**
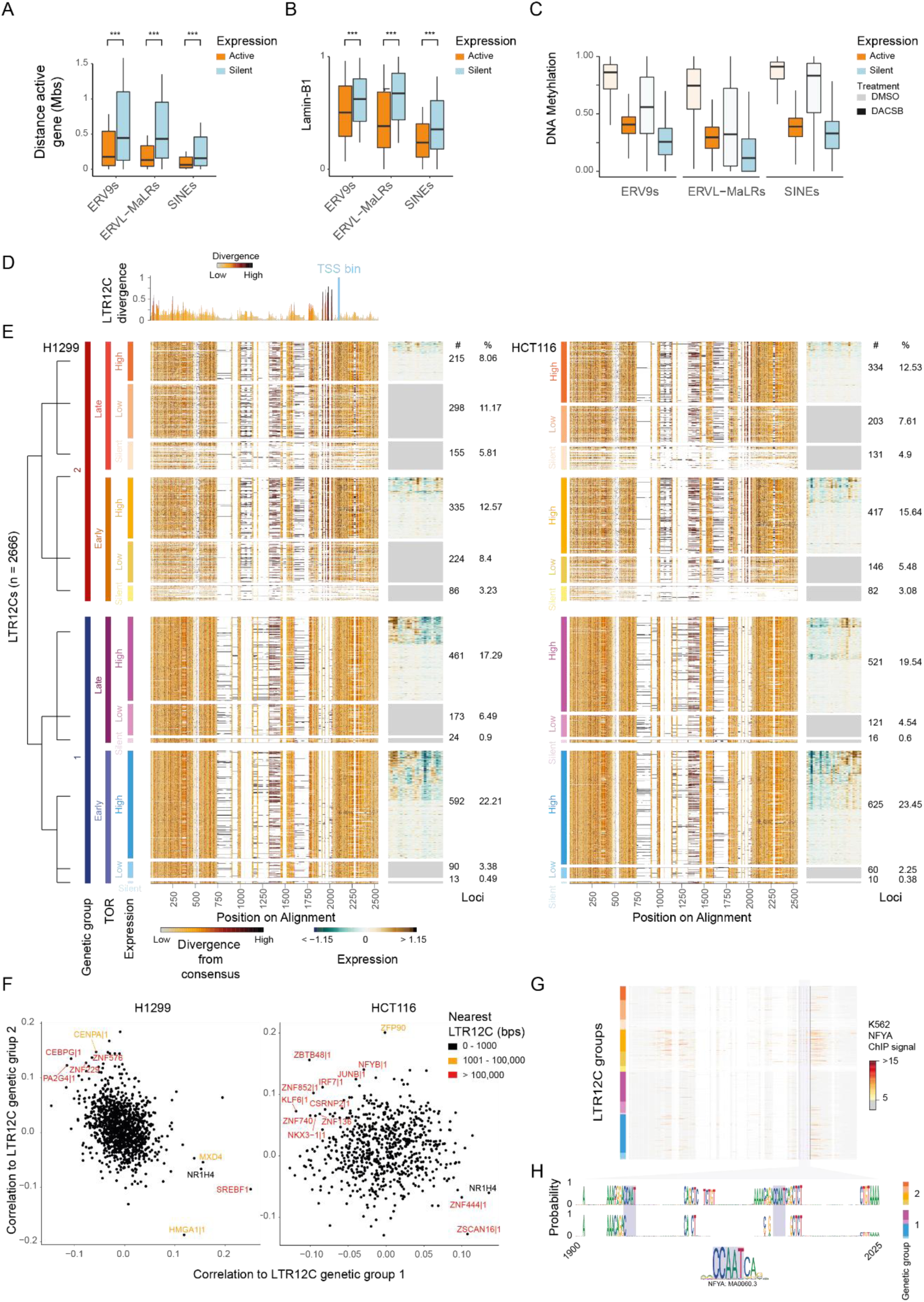
Sequence composition and genomic context explain TE transcriptional activation and variability. A) Boxplot showing the distance to the nearest identified genic TSS for active (>10 TSS UMIs, orange) and silent (0 TSS and non-TSS UMIs, blue) TE loci grouped by TE class/superfamily membership in H1299 cells. Boxplot whiskers extend 0.5 times the interquartile range from the 25th and 75th percentiles and loci more than 2 mega bases away are not shown. B) Lamina-association of active (orange) and silent (blue) TE loci grouped by TE class/superfamily in H1299 cells. C) Distribution of average methylation for active (orange) and silent (blue) TE loci before (light colors) and after treatment (dark colors) grouped TE class/superfamily membership in H1299 cells. D) Rolling mean (five nucleotides) of the sequence divergence between two genetic subgroups of highly expressed LTR12C elements. Colors indicate the degree of divergence. E) Multiple sequence alignment (left) and TE metacell expression (right) for 2666 LTR12C loci hierarchically grouped by genetic sequence (clusters #1 and #2), time of replication (early and late), and transcriptional activity (silent (0 UMIs), low (less than 10 TSS UMIs) and high (at least 11 TSS UMIs)). Loci (rows) within a group are arranged according to K-means clustering of TE metacell footprints. Numbers to the right indicate the number of loci and their percentage from total for each group. Colors either represent the position-wise genetic divergence from the consensus or the TE metacell footprint. Alignment is identical for H1299 (left panel) and HCT116 (right panel) cells, but the order may differ according to cell-line specific expression dynamics. F) Spearman correlation of TF expression levels to summarized LTR12C genetic group 1 and 2 loci counts (all downsampled). Color of annotated TFs represents distance to nearest highly expressed LTR12C element. G) K562 ChIP-seq signal for NFYA on top of the LTR12C multiple sequence alignment. Individual loci are grouped according to the H1299 data shown in E. H) Consensus sequence logos from position 1900 to 2025 on the alignment for group 1 and 2 loci. Putative NFY motif sites are highlighted.

In addition to genomic context and epigenomic markup, sequence divergence between TEs within the same family may be predictive of de-repression trends^14^. To test this, we clustered 2666 (excluding 66 loci on chromosome Y) sequences from the LTR12C family according to their sequence, deriving two distinct clusters varying by hyper-variable regions at multiple positions and in particular close to the identified TSS (**Fig 4D**). We then hierarchically classified elements in each of the two clusters according to their time of replication and analyzed the overall expression in each group, as well as the distribution of expression over TE metacells (**Fig 4E**). The data showed substantially strong expression in genetic cluster #2 compared to #1 (p << 0.01 in both cell lines, Wilcox). Within the genetic clusters, we still observed a strong correlation between time of replication and activity patterns (p<< 0.01 in both cell lines, Wilcox), highlighting how broad genomic context and sequence composition additively contribute to the de-repression propensity within a TE family.

Interestingly, overall expression intensities of both clusters could be disconcordant and even inversely correlated at the single-cell level (**Fig S7C**), suggesting sequence variations in cis may be interpreted by a cell’s TF arsenal resulting in differential transcriptional activation. In search of such putative regulators, we identified several candidate TFs with cluster-specific correlations, including multiple zinc finger proteins and other genes with reported function in retrovirus biology (e.g. IRF7^7^) (**Fig 4F**). To our surprise, NFY, a known activator of LTR12C expression^20^, was positively associated with expression of cluster 2 loci while lacking association with the other group in HCT116 cells. We hypothesized that partial regulation by NFY of a subset of loci should be reflected by differential binding affinities of the trimeric NFY complex. Indeed, NFYA ChIP-seq signal^21^ from two cancer cell lines showed specific coverage for cluster 2 loci (p << 0.01, Wilcox), especially those in early replicating compartments (**Fig 4G and Fig7D**). The lack of binding affinity towards cluster 1 loci coincided with a specific and highly localized degeneration of NFY DNA binding sites, directly upstream of the TSS (**Fig 4H**). Together, the data suggest that cluster #2 TEs are de-repressed independently of NFYA availability, possibly given permissive epigenomic contexts that are receptive to more general activation machinery, while Cluster #1 TEs are dependent on NFYA for driving derepression in less favorable epigenomic contexts.

## DISCUSSION

We performed 5’ single-cell transcriptional profiling to *de novo* identify TSS activity in epigenetically de-repressed cancer cells. Using a computational approach that leverages the high genomic copy-number and sequence conservation of TE families, we robustly pinpointed the exact sites and degree of transcription initiation in thousands of loci representing the major transposon classes. This uncovered a rich and dynamic landscape of TE cell-to-cell variation in supposedly homogenous cancer populations. We observed different overall transcriptional dynamics for the four major TE classes. SINE retrotransposons largely showed a pattern of expression consistent with a more sporadic and spurious de-repression, although globally correlated with cellular proliferation. Specific families of DNA transposons and LINEs were linked to a presumed pre-apoptotic phenotype, suggesting broad-scale loss of genomic control. On the other hand and consistent with a previous report^22^, we observed structured activity of LTR retrotransposons, which was linked to specific stages of the cell cycle and additional genic pathways. Coordinated activity of multiple loci from the same TE family was observed systematically, but in some cases, and contrary to our expectations, loci of the same LTR family could be clustered into modules showing distinct combinatorial expression patterns. This data shows that even in cells with largely perturbed epigenetic integrity, TE derepression is still regulated by trans-factors (through cell cycle fluctuation or more stable transcriptional states).

As exemplified for members of the massively de-repressed LTR12C TE family, intra-family expression variability is associated with at least three layers of regulation. First, activated loci rely on a favorable broad genomic context and proximity to activator machinery at constitutively active TSSs. Second, de-repression involve change in local epigenetic composition (in particular DNA methylation) of the TE TSSs. Finally, specific sequence characteristics within the TE family consensus module are linked with locus-specific regulation, ensuring or preventing effective recruitment of the activatory trans-machinery, as we have exemplified for NFY. Single cell RNA-seq can uncover the combinatorics of these layers and how they affect TE regulation in general, improving our understanding of the previously described cell-type and disease-specific TE expression landscapes^23–25^.

Understanding the genomic rules of TE de-repression is particularly important in the context of epigenetic therapy, where a subset of ERVs is reported to mimic a viral-like interferon response and to serve as a rich source of potentially immunogenic neoantigens^7,8,14^. We found the expression of several LTR families to be correlated with overall interferon type-I gene expression at single cell resolution. Such variability inherently represents differential response to treatment, motivating further studies on the interplay between TE activity, intra-cellular regulation of immune response, and the efficacy of epigenetic and immunotherapy. Epigenetic relaxation is also observed during massive genome-wide DNA demethylation during embryogenesis, germline development, or when cells progressively lose their canonical DNA methylation in ageing^26–28^. The pervasive regulation of TE dynamics we observed in this study suggest the possibility that a large number of un-appreciated hotspots of genomic activity can become activated in such processes. Incorporating single cell TE activity profiles into models of epigenetic control and gene regulation will provide a powerful tool for understanding TE sub-functionalization toward developmental enhancers and alternative promoters, as well as in the context of aberrant de-repression leading to disease.

## DATA ACCESS

Raw and processed data were deposited in the gene expression omnibus repository under accession GSE121309.

## MATERIAL AND METHODS

### Cell culture and treatment

HCT116 parental colorectal cell lines (HD PAR-033) were obtained from Horizon Discovery Ltd, Cambridge, UK and NCI-H1299 non-small cell lung cancer cells were provided by the courtesy of Prof. Moshe Oren. H1299 cells were cultured in RPMI 1640 (Gibco 21875034I) and HCT116 cells were grown in heat-inactivated McCoy’s 5A medium (Biological Industries, Israel; 01-075-1A), both supplemented with 10% Fetal Bovine Serum (Gibco FBS 10270-106), 0.4% Penicillin-Streptomycin (Biological Industries, Israel; 01-031-1C) and 1% L-glutamine (Biological Industries, Israel; 01-020-1). Conditioned media were filtered through a 0.22 microns filter (Corning, 430769) prior to culture. Cells were split at a ratio of 1:10 every 2–3 days using 0.05% trypsin-EDTA solution C (Biological Industries, Israel; 03-053-1B). Treatment with the DNA hypomethylating agent 5-Aza-2′-deoxycytidine (Sigma-Aldrich, A3656; Lot #SLBS4457) and HDAC inhibitor SB939 (Cayman Chemical, 10443; Lot# 0435336-34) was performed as previously described^14^. To prepare single-cell suspensions, cells were processed according to the instructions of the 10x Genomics Single Cells Protocols Cell Preparation Guide (10x Genomics, CG00053 Rev C). About 4000 cells per cell line and treatment condition were used as input for further processing.

### 5’ single-cell RNA sequencing

Libraries were generated using the Chromium Single Cell 5’ Library & Gel Bead (10x Genomics, PN-1000014) according to the manufacturer’s instructions and sequenced on the Illumina NextSeq 550 using the High Output v2 kit (Illumina, TG-160-2004).

### Curation and binning of gene and TE models

Gene models were based on Gencode release 28 (GRCh38.p12) obtained from (https://www.gencodegenes.org/releases/). Exon coordinates of all transcript isoforms were reduced to a single set per gene symbol using the reduce function of the R package GenomicRanges. To minimize mapping ambiguity for exons shared between different genes, we then created a set of non-overlapping exon coordinates with the disjoin function of the same package and retained all gene symbols for shared intervals. After elongating the 5’ end of exons containing annotated TSSs to 100bps or to the next exon, we binned the resulting coordinates into 20bp intervals.

We downloaded genomic TE alignments generated by RepeatMasker open-4.0.5^29^ with the repeat library 20140131 from (http://www.repeatmasker.org/species/hg.html). The obtained alignments were used to map genomic TE sequences onto their relative position on the corresponding consensus model at 20bp resolution (**Fig S8**). Rare TE sequence polymorphisms not aligning to the corresponding consensus model were assigned to the closest preceding consensus alignment. In case resulting 20bp genomic TE bins ambiguously mapped to more than one TE family, only the family with highest alignment score was retained. To minimize misassignment of genic UMIs onto TE loci, we finally excluded TE bins located within genic exons (except those overlapping any genic reference TSS) to generate the final TE coordinate set.

### Data processing

Read pairs missing the constant part of the template switch oligo (defined by a hamming distance to TTTCTTATATGGG greater than 4) were excluded before subsequent processing with the cell-ranger analysis software version 2.1.0. Alignment was performed against the human GRCh38-1.2.0 reference using non-default Cell Ranger parameters (chemistry = SC5P-PE; r1_length = 100; r2_length = 50) and the alignment-specific parameter (--outFilterMultimapScoreRange: 2) for improved mapping specificity. UMI tools^30^ (--per-cell; --extract-umi-method = tag; --buffer-whole-contig; --cell-tag = CR; --umi-tag = UR; --paired) was used to remove duplicates. Barcodes were classified as cell-associated using the standard Cell Ranger (2.1.0) procedure. Identified cell barcodes with low mitochondrial (stripped nuclei) or ribosomal counts were further excluded from this set (**Fig S1A**). Coordinates of the first position of read1 in properly and uniquely aligned read pairs were tested for overlaps with binned exon or TE intervals and used for downstream analysis.

### Alignment benchmark

The impact of throwing multi-mapping reads before TE quantification was assessed by counting the number of unique and multi-mapping reads (**Fig S2A and B**). In case multiple alignments (up to 10 distinct genomic positions) mapped to the same TE family, the read was assigned to the family with maximal count (ties were resolved randomly). Family-wise mappability score was then defined as the percentage of uniquely mapped from all UMIs (including multi-mappers).

Pairwise hamming distances between reads mapping to the same TE family were calculated by first extracting first mates of read pairs aligning to LTR12C loci on the plus strand of chromosome 1. To minimize the impact of frameshifts, only mates mapping to the maximally covered position in each locus were retained and the number of mates was capped at 100 per locus. The resulting 2425 × 2425 comparisons representing mates mapping to 65 expressed LTR12C loci are shown in **Fig S3D**.

Read error simulations were performed using the mutateReads function of our Reputils R package (source code can be found here: https://tanaylab.bitbucket.io/Reputils). Briefly, reference sequences of one million sampled TE alignments were extracted, randomly mutated using 0.25, 0.5, 1, 2, 4, and 8% global error rates, and re-aligned against the genome using mapping parameters as before.

### TSS mapping

UMI counts were aggregated for all cells per TE family or gene bin. To account for TE inter- and intrafamily bin copy-number and size variations, TE aggregate bin UMI counts were normalized by a bin’s genomic copy-number and median width, and regularized by the families’ median bin copy-number (**Fig S8**). Genic bins with a minimal UMI to total gene expression ratio of 0.1 and at least 10 raw UMIs were marked as putative TSSs and combined if less than 100bp apart. For TE bins, we applied more stringent TSS selection criteria due to the lack of reference annotation and higher expected background signal. Specifically, we calculated the UMI enrichment over the neighboring 8 bins (with an offset of 1 in both directions) and assessed statistical significance based on the Poisson distribution. TE bins with at least 20 raw UMIs, fold-change > 3, and FDR corrected p-value < 0.01 were selected as putative TSS. While both genic TSS and non-TSS UMIs were retained (distinguished by an |# TSS suffix), only UMI signal from individual TE loci corresponding to the identified TSS bin were kept before summarizing molecule counts in a UMI matrix U = [u_fi_] on features f and cells i. For gene-level analyses, we focused on protein-coding genes and excluded non-TSS UMI counts in case a gene was associated with an enriched TSS. This exclusion was done to minimize the impact of UMI signal from de-repressed intronic TE loci bleeding into reference gene models^14^.

### Metacell analysis

#### Transposable elements

We used the MetaCell package^31^ version 0.3.39 for TE feature selection, grouping of cells, and to calculate gene enrichment along those groups, with the following adaptations. No preliminary cell filtering was applied to TE UMI counts. Markers to model cell-to-cell similarity were selected based on a normalized variance threshold above 0.1, a total UMI count of at least 10 molecules, and second highest UMI count of 2 after downsampling to 401 (H1299) and 370 (HCT116) TE TSS UMIs. After constructing an initial similarity graph with a K = 100 parameter, we sampled 500 metacell covers with a minimal cell size of 50, each organizing 75% of cells into coherent groups in the graph. The resulting pairwise co-clustering of all single cells was used as input to construct the final similarity graph and robust metacell covers were inferred using default parameters.

#### Cell cycle

Cell cycle-related genes were defined based on their correlation to one of the marker genes PCNA, E2F1, TOP2A, MKI67, UBE2S, ATAD2, ARL6IP1, AURKA, or RRM2 (r >= 0.14 to 0.25). Clustering correlated genes based on downsampled single-cell UMI count matrices, was followed by manual selection of gene clusters that consisted of cohesive cell cycle gene modules. This procedure was repeated for all conditions using four different down-sampling depths (n=10000 to 5000 genic UMIs) to finally select 63 - 119 cell cycle regulated genes for further analysis. Metacell analysis of a UMI matrix consisting of these genes alone (aiming at larger metacell sizes by setting min_mc_size = 100) was then performed. The resulting cell cycle metacells were arranged according to their estimated phase (From G0/1 to mitotic exit) in a supervised fashion based on the expression of marker genes (**Fig S6A**), organizing all cells along a putative cell cycle trajectory.

### Cell cycle effect on TE expression

2D projection of single cells and metacells according to the cell cycle model was performed using the standard Metacell projection algorithm setting T_edge to 0.05^31^. To plot cell cycle progression of individual TE modules, we used the grouping of cells according to the genic cell cycle module. Downsampled expression levels of TE modules were summarized per cell and their enrichment was compared to the module’s mean expression intensity across all cells. The resulting enrichment score was visualized on top of the cell cycle projection or log2 transformed (a pseudocount of 1 was added) before applying non-parametric local regression (locally weighted smoothing) across cells grouped by their cell cycle metacell stage. This approach provided adequate power for testing correlation of TEs with different cell cycle phases, and we therefore did not aim at more continuous modelling of the transcriptional dynamics as suggested by more direct modelling strategies^32,33^.

### Time of replication, lamin association, and NFY ChIP-seq

Repli-seq and DAM-ID normalized coverage files were obtained from the 4D nucleus^34^ data portal (https://data.4dnucleome.org/) for HCT116 and H1-hESC cells, respectively. Replicates were averaged and resulting coverage values were percent rank and log2 transformed. NFYA fold change over control signal from two replicates in K562 cells was downloaded from the ENCODE^21^ data portal and percent rank transformed.

### Multiple sequence alignment and genetic divergence

MAFFT^35^ version 7.394 was used to align sequences with a gap opening penalty and offset of 1.1 and 0, respectively. Positions with a gap frequency greater than 99% on the alignment were removed for visualization. Distances between TE loci were computed using the dist.dna function of the R package ape using the indelblock substitution model. Ward’s linkage was used to group loci. A locus’ position-wise divergence from the consensus was defined as 1 – that bases’ consensus frequency (including gaps as a fifth base). Group-wise sequence divergence was calculated based on the maximal delta in base frequency per position.

### Selection of IFN-I genes

All annotated IFN-I genes from the interferome^36^ database were downloaded and further filtered for genes that were consistently upregulated with a minimal average fold-change of 2 across at least 2 independent studies. We compared expression between untreated and treated cells for the remaining 1220 putative IFN-I regulated genes and defined a final set of 164 (H1299) and 127 (HCT116) genes with increased (log2 fold-change larger than 2) expression following treatment.

### DNA methylation analysis

DNA methylation values for untreated and treated H1299 cells were downloaded from the gene expression omnibus repository under accession GSE81322. Hg19 CpG coordinates were lifted to hg38 using the liftOver function from the Genomic Ranges R package.

### Statistical analysis

All statistical analyses were performed using the R statistical environment, version 3.4.2. Unless stated otherwise, box plot center lines indicate data medians, box limits indicate the 25th and 75th percentiles, whiskers extend 1.5 times the interquartile range from the 25th and 75th percentiles, and outliers are not shown. For group-wise comparison of two distributions from different samples or treatments, the two-tailed nonparametric Wilcoxon and Mann-Whitney test was used. Unless stated otherwise, p-values < 0.05 were considered statistically significant and significance levels are depicted as follows: * P < 0.05, ** P < 0.005, *** P < 0.0005. The Benjamini and Hochberg method was used to correct for multiple hypothesis testing.

## Code availability

Analysis scripts and processed data files are available under (https://bitbucket.org/tanaylab/brocks_et_al_nat_com_2019_epitherapy_scrna/src/default/).

## ACKNOWLEDGEMENTS

We thank L. Velten and C. Plass for helpful feedback and discussions and Z. Meir and A. Sebe-Pedros for critical proofreading of the manuscript. We are also grateful for the support of H. Keren-Shaul and other members of the Advanced Sequencing Technologies facility at the Weizmann Institute of Science. D.B. is supported by an EMBO long-term fellowship (ALTF 900-2017). Research was funded by the European Research Council (grant scAssembly), by the flight attendant medical research institute and the ISF I-core program. We thank B. van Steensel and D. Gilbert for providing access to lamin association and time of replication data generated as part of the 4DN NIH consortium.

