## Supplemental figures for "Single cell analysis reveals dynamics of transposable element transcription following epigenetic de-repression"

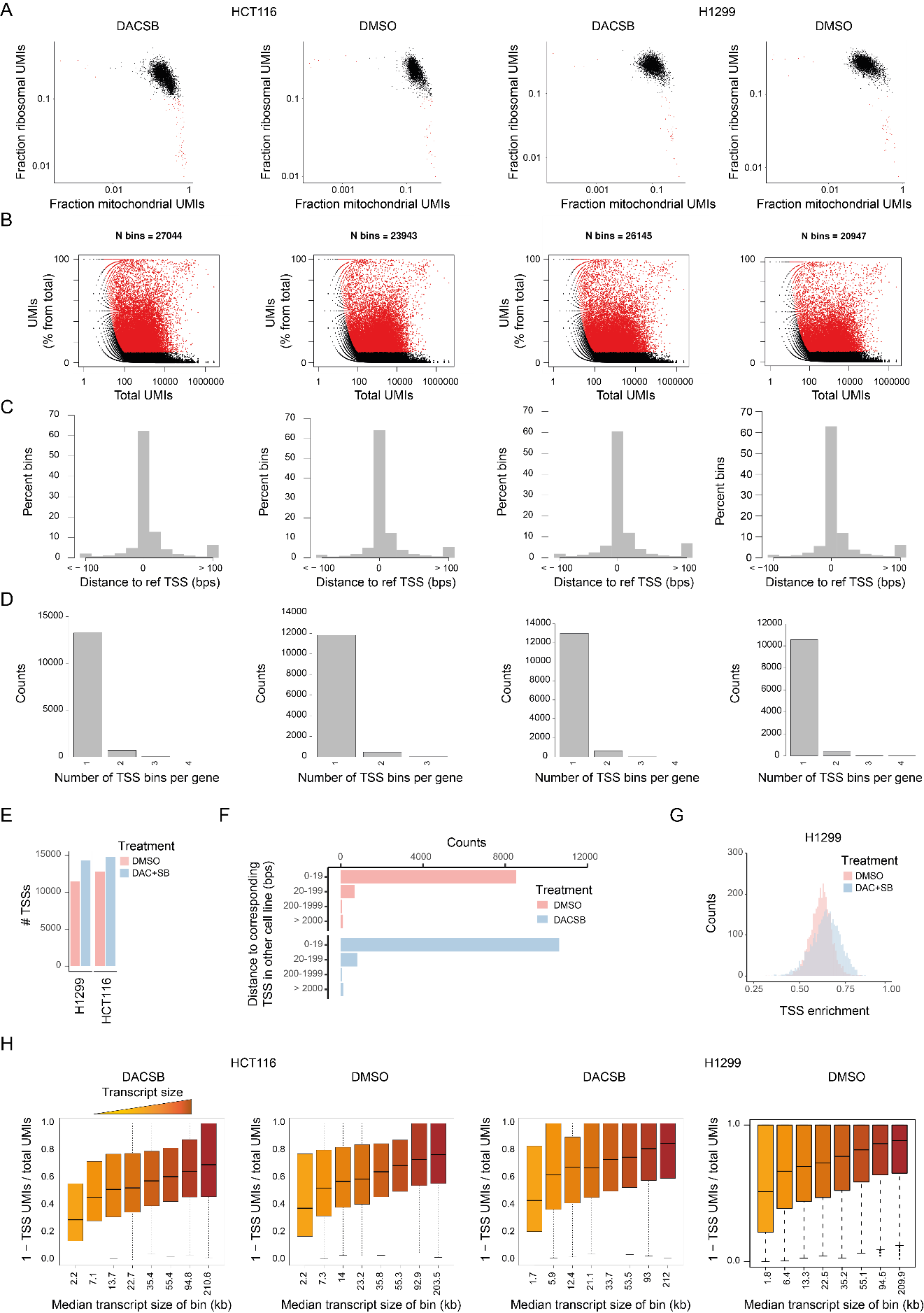

**Supplementary Figure 1**

A) Fraction of UMIs mapping to ribosomal versus mitochondrial genes (characterized by an RP[LS] and MT- prefix in their gene name, respectively). Cells highlighted in red were excluded from downstream analyses.
B) Fraction of bin UMI coverage as a function of total coverage per gene. Putative TSS bins are highlighted in red.
C) Histogram of the genomic distance of selected bins to the nearest reference TSSs.
D) Counts of distinct putative TSS bins per gene model.
E) Final number of identified distinct TSS bins in untreated (red) and treated (blue) cells.
F) Counts of distances to nearest corresponding TSS in the other cell line for genes expressed in both cell lines.
G) Single-cell distribution of TSS enrichment (proportion of UMIs mapping to identified TSSs) before (red) and after (blue) treatment shown for H1299 cells.
H) Gene-wise TSS enrichment stratified by groups of increasing spliced transcript size based on cell aggregate counts

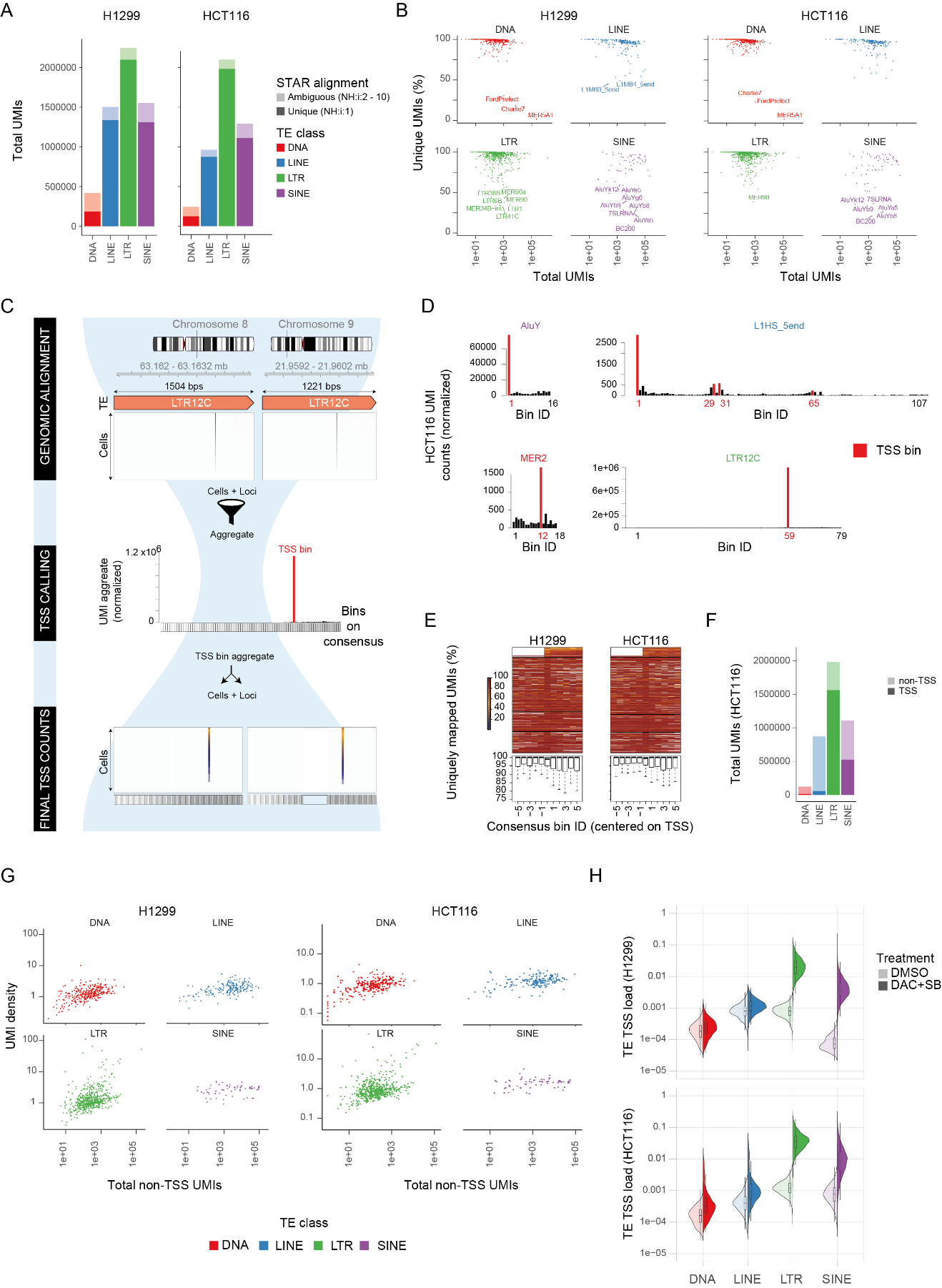

**Supplementary Figure 2**

A) Number of unique and multi-mapping UMIs per TE class.
B) Per family UMI mappability grouped by TE class. Families below 50% unique UMIs are highlighted.
C) Strategy to identify putative TE TSSs. UMIs aligning unambiguously to single genomic TE copies are aggregated across all cells and all loci of a given family. UMI binning over the consensus model is performed at 20bp resolution, as exemplified for two LTR12C copies (top) (see also **Fig S8**). Consensus TSS bins are selected by their UMI enrichment compared to neighboring consensus bins (middle). UMI signal from identified TSS bins only is then projected back to individual loci and single cells. Downstream analyses are performed on signatures of transcription for individual TE loci and single cells (bottom).
D) Distribution of normalized UMI counts along the consensus model of selected (highest overall expression in H1299) representatives from each TE class in HCT116 cells. Putative TSS bins are highlighted in red.
E) Mappability score defined as the proportion of uniquely mapped UMIs from the total (including UMIs with up to 10 distinct genomic positions) per family (top) and summarized (bottom). Families are arranged as in **Fig 1E**.
F) Number of TSS and non-TSS UMIs mapping to different TE classes in treated HCT116 cells.
G) Family-wise UMI density (defined as non-TSS UMIs per genomic kilobase) over total number of non-TSS UMIs.
H) TE load distribution across cells based exclusively on TE TSS UMIs.

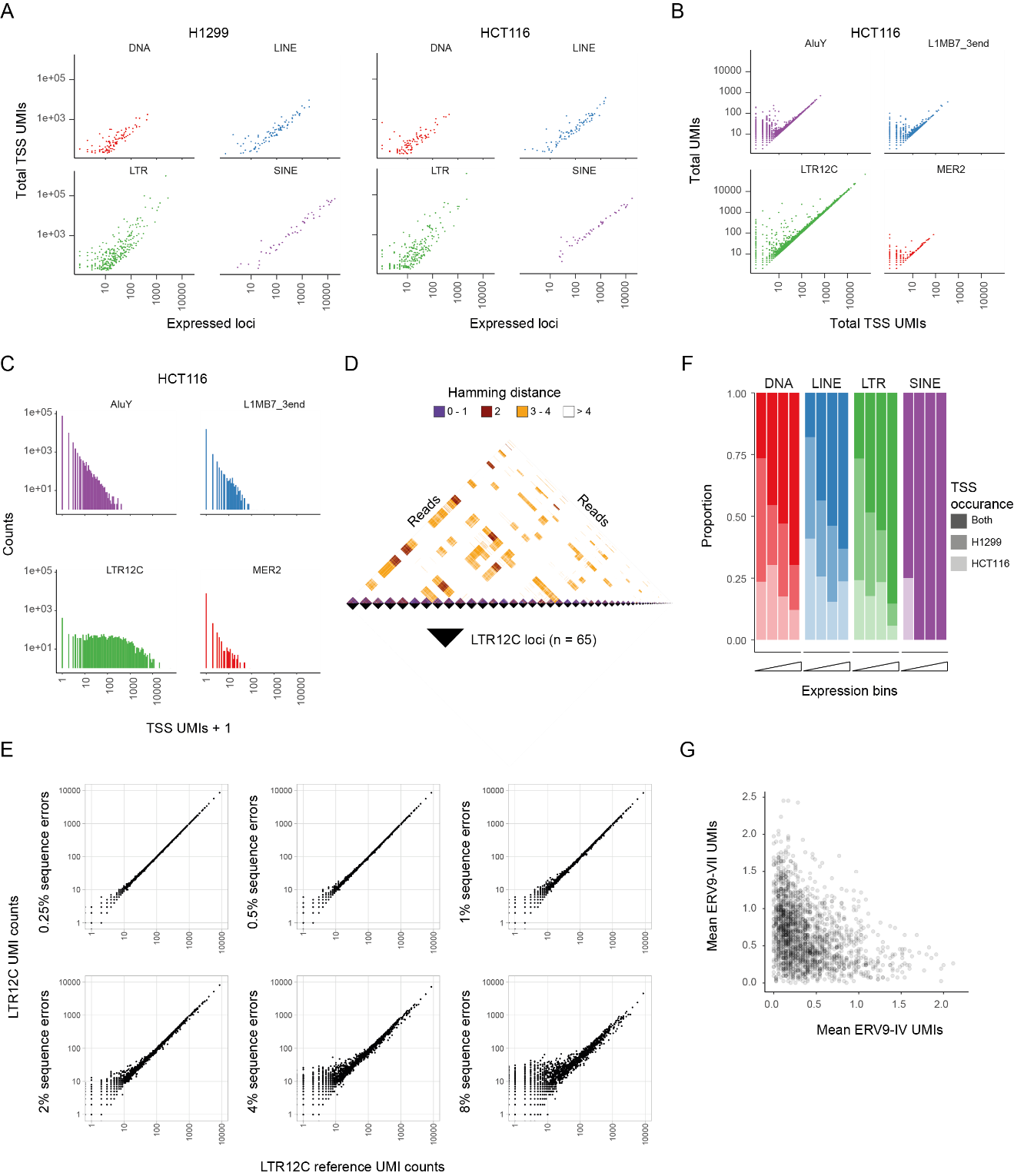

**Supplementary Figure 3**

A) Per family total number of TSS UMIs by the number of active loci (at least 1 UMI).
B) Per locus total number of UMIs as a function of TSS only UMIs for representative TE families (highest expression in HCT116 per class).
C) Distribution of total UMIs per locus for representative TE families.
D) Pairwise hamming distances between first mates mapping to LTR12Cs on chromosome 1 in treated H1299 cells. Mates are grouped by their originating LTR12C locus.
E) Comparison of reference LTR12C UMI aggregate counts against counts obtained from reads with increasing levels of base substitution errors.
F) Proportion of shared and cell line specific TSS bins for families with putative TSS activity in at least one cell line, stratified by TE class and total expression.
G) Per cell summarized mean expression of ERV9-VII vs ERV9-IV TE modules based on downsampled UMI counts in H1299 cells.

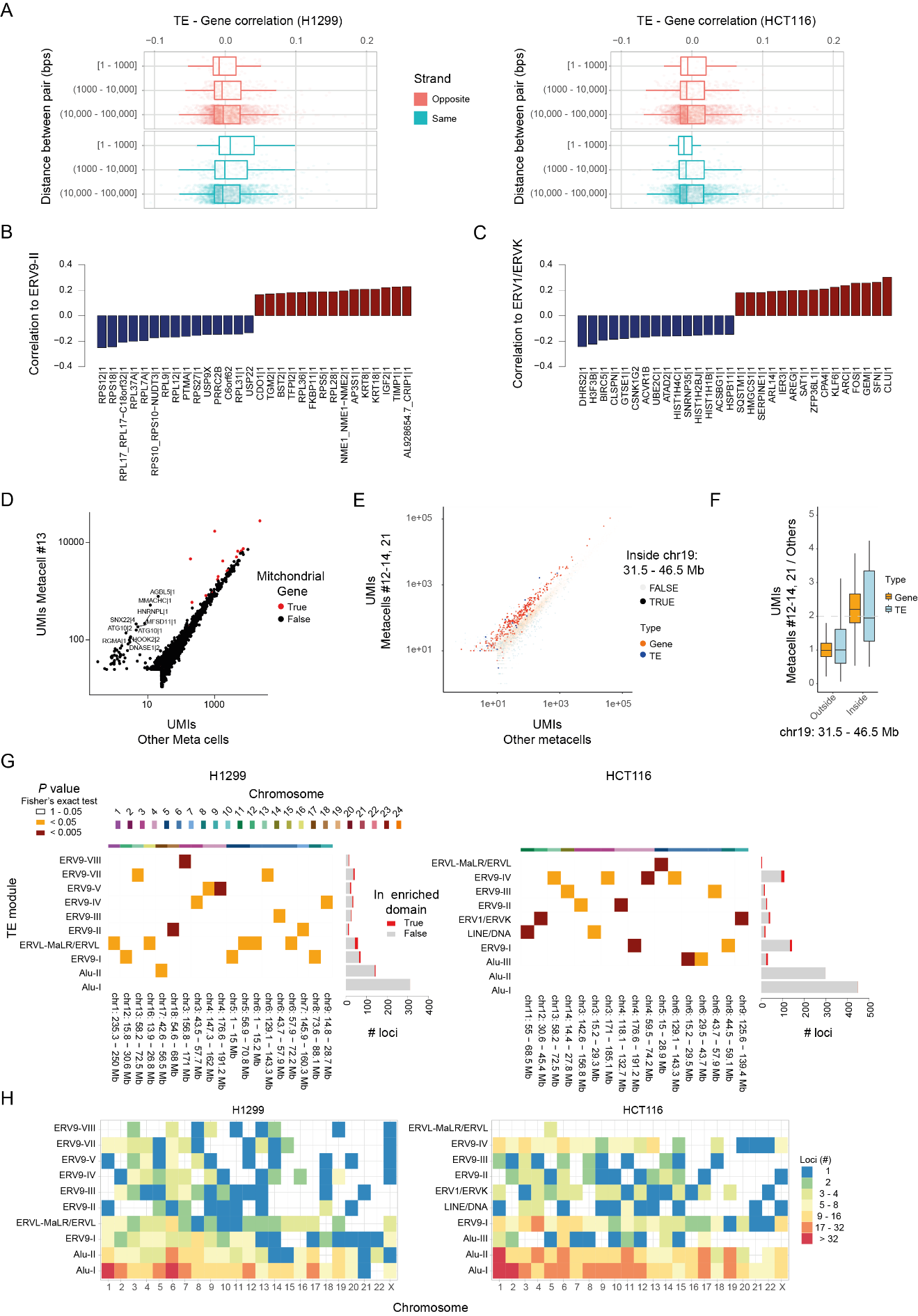

**Supplementary Figure 4**

A) TE vs gene pairwise expression correlation grouped by base pair distance between their TSSs. Only Pearson correlation between -0.1 and 0.2 is shown.
B) Top (anti)-correlated genes to the ERV9-II module in H1299 cells.
C) Correlation of selected genes to the HCT116 ERV1/ERVK module.
D) Summarized gene expression in HCT116 metacell #13 vs all others.
E) Genic and TE expression in H1299 metacells #12-14 and 21 vs all others. Feature located within and outside chromosomal domain 31.5 – 46.5 on chromosome 19 are marked by dark and light coloring, respectively.
F) Fold-change TE and genic expression between H1299 metacells #12-14 and 21 vs all others. Vertical dashed line indicates a fold-change of 2.
G) Enrichment of 15 megabase chromosomal domains for TE loci of the indicated modules in H1299 and HCT116 cells and number of loci per module mapping to statistically enriched domains.
H) Number of loci mapping to different chromosomes (columns) per TE module (rows).

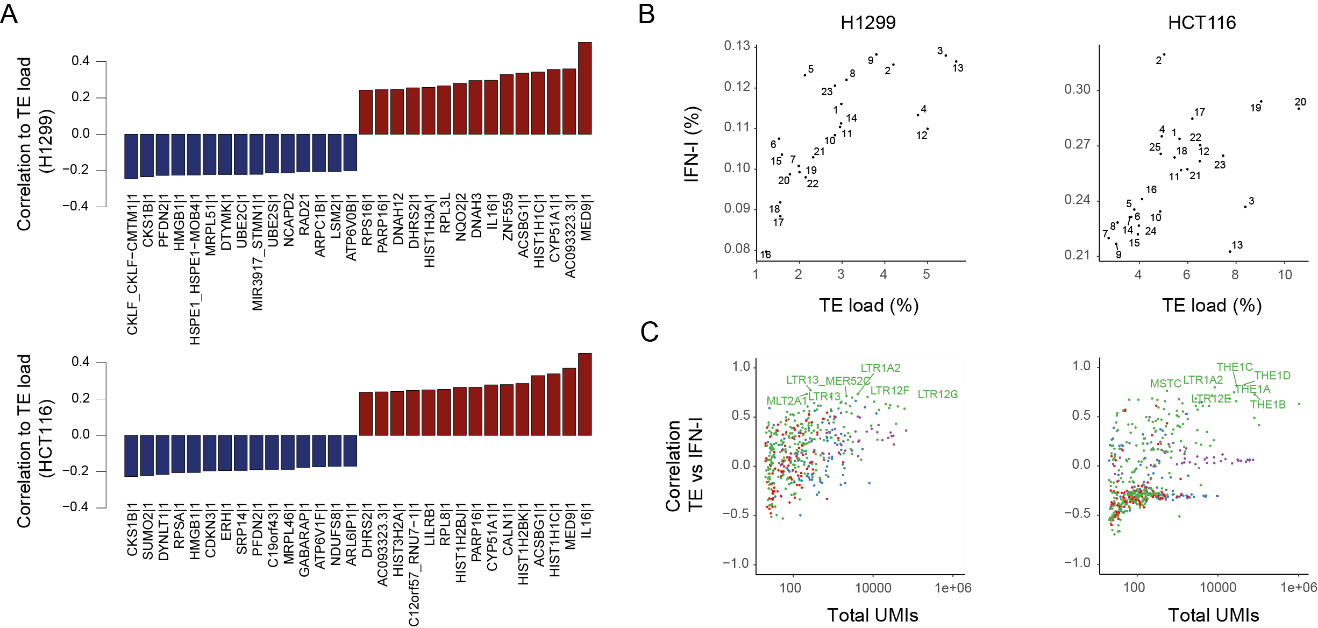

**Supplementary Figure 5**

A) Top (anti)-correlated genes to TE load in H1299 and HCT116 cells.
B) Percentages of IFN-I intensity vs TE load based on aggregated TE metacell UMI counts in H1299 and HCT116 cells. Numbers indicate TE metacell ID.
C) Pearson correlation of TE family load vs IFN-I intensity calculated at the metacell level. Families with a correlation >0.7 are labeled. Colors indicate TE class.

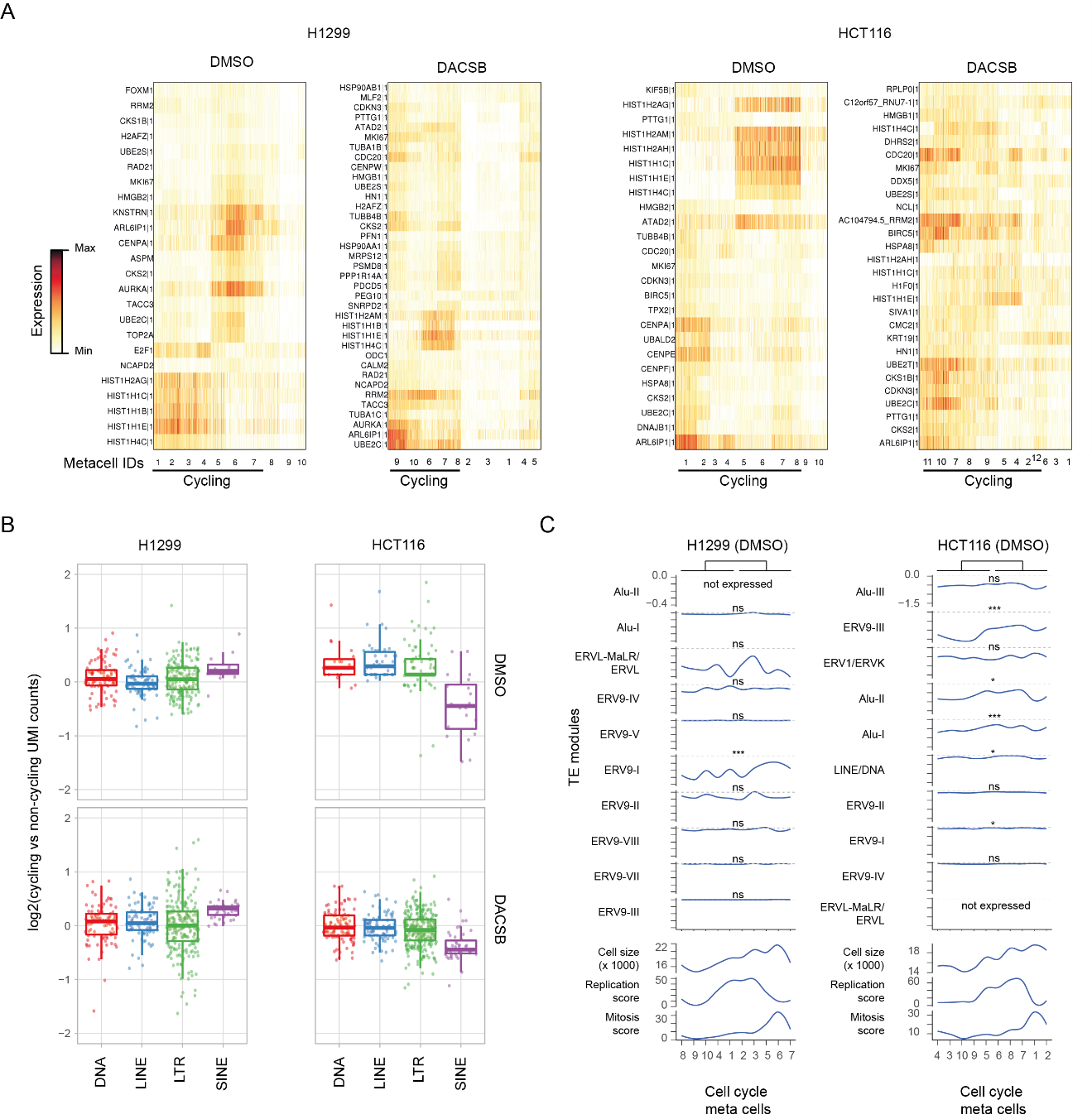

**Supplementary Figure 6**

A) Single-cell expression matrix of the indicated conditions. Each column represents individual cells and bottom numbers indicate cell cycle metacell IDs. Rows represent genes with differential metacell footprint.
B) Expression fold-change of TE families (dots) between cycling and non-cycling cells. UMI counts were summarized per TE family across all cycling or non-cycling cells and scaled by dividing by the total number of molecules in the smaller (less molecules) condition.
C) Raw UMI enrichment of TE modules identified after treatment along the cell cycle trajectory of untreated cells (as in **Fig 3F**). Significance is based on Wilcoxon rank-sum test comparing cells from the first five metacells on the inferred cell cycle trajectory (#1, 4, 8-10 and #3-5, 9-10 in H1299 and HCT116, respectively) against the latter five.

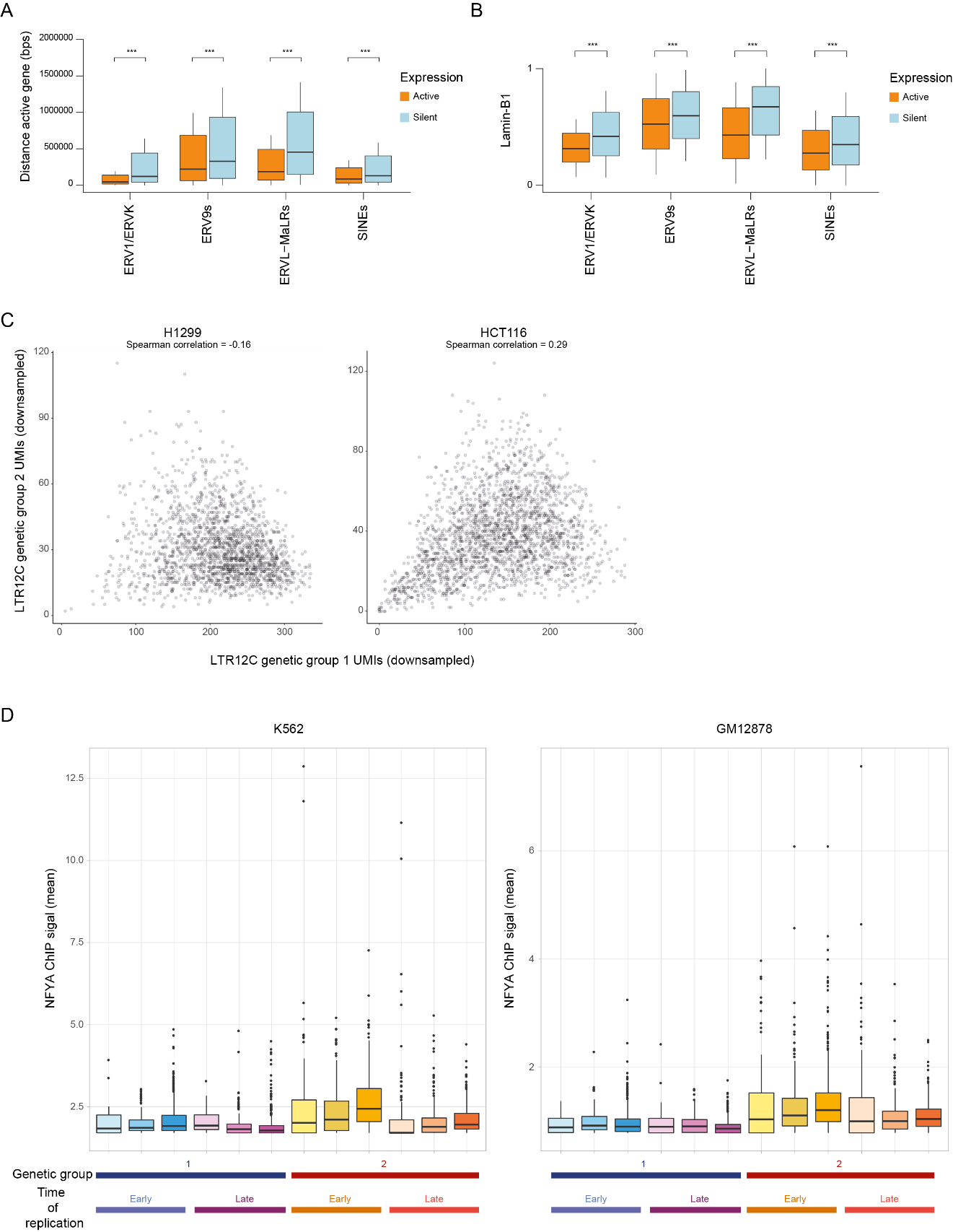

**Supplementary Figure 7**

A) Boxplot showing the distance to the nearest identified genic TSS for active (>10 TSS UMIs, orange) and silent (0 TSS and non-TSS UMIs, blue) TE loci grouped by TE class/superfamily membership in HCT116 cells. Boxplot whiskers extend 0.5 times the interquartile range from the 25th and 75th percentiles and loci more than 2 mega bases away are not shown.
B) Lamina-association of active (orange) and silent (blue) TE loci grouped by TE class/superfamily in HCT116 cells.
C) Per cell summarized total expression of genetic group 1 and 2 loci based on downsampled UMI counts.
D) Distribution of mean NFYA ChIP signal per locus grouped by genetic cluster, time of replication, and expression intensity in treated H1299 cells (as in **Fig 4E**).

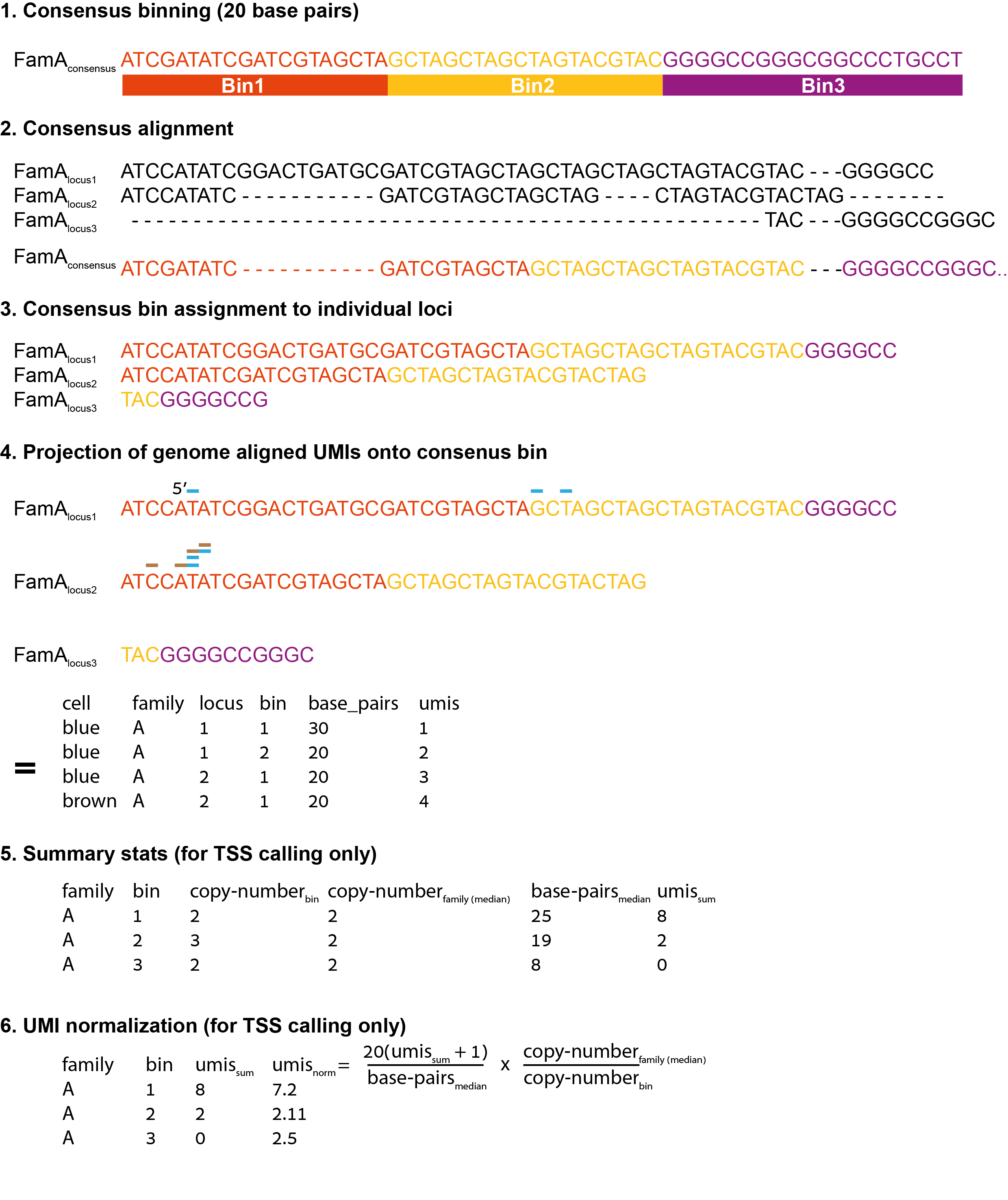

**Supplementary Figure 8**

Workflow to assign genome mapped reads onto consensus bins for TSS calling. (1) TE consensus sequences were binned at 20 base pair resolution. (2) Individual genomic TE loci were then pairwise aligned against their corresponding consensus model. (3) Each position of genomic TE loci was matched to the corresponding 20 base pair bin on the consensus based on the resulting alignment. Rare sequence polymorphisms not represented in the consensus model (for example TAG 3’ end of locus 2) were assigned to the nearest preceding bin ID. (4) Uniquely mapped genomic read alignments were then grouped by TE family and bin ID, (5) and finally normalized to account for variable bin copy-number and size distributions (due to genomic indels) before TSS identification.
